# MCBO: Mammalian Cell Bioprocessing Ontology, A Hub-and-Spoke, IOF-Anchored Application Ontology

**DOI:** 10.64898/2026.01.05.697007

**Authors:** Kimberly Robasky, James Morrissey, Markus Riedl, Andreas Dräger, Nicole Borth, Michael J. Betenbaugh, Nathan E. Lewis

**Author notes:** Corresponding author. (K. Robasky); (N. E. Lewis). These authors contributed equally (Kimberly Robasky, James Morrissey, Markus Riedl, Andreas Dräger, Nathan E. Lewis).

## Abstract

Mammalian cell-based biopharmaceutical manufacturing generates vast, heterogeneous datasets that remain fragmented due to the lack of a standardized metadata framework. A key challenge in biologics manufacturing is linking bioreactor conditions, cell line characteristics and recombinant product production. A datahub for mammalian cell bioprocessing, integrated by semantic technologies, will serve as a tool to understand and query the connections between these complex datasets. While existing ontologies cover general biological and experimental concepts, they often lack the operational specificity required to harmonize bioreactor conditions, cell line engineering, and product quality metrics. To address this specific gap, we present the Mammalian Cell Bioprocessing Ontology (MCBO), a hub-and-spoke application ontology built on Basic Formal Ontology (BFO) foundations and anchored to the Industrial Ontology Foundry (IOF) Core. MCBO formalizes the process-participant-quality modeling pattern, enabling precise tracking of culture environmental conditions as qualities of the physical culture system. We demonstrate the utility of MCBO through a central datahub populated with 723 curated cell culture process instances and 325 unique bioprocess samples from published studies. The framework is validated against eight competency questions implemented via SPARQL, demonstrating efficient cross-study querying of culture optimization, cell line engineering, and multi-omics integration. By providing a stable, schema-independent substrate for data harmonization, MCBO enables AI agent-powered, human-in-the-loop workflows and facilitates LLM-assisted extraction of structured metadata from legacy records. MCBO is open-source and designed for deployment behind institutional firewalls to support interoperable biomanufacturing intelligence while maintaining intellectual property sensitivity. MCBO is supported by the International Biomanufacturing Network (IBioNe), which aims to accelerate discoveries and developments by providing a network of biomanufacturing training and workforce development to educate the next generation of biomanufacturing experts. MCBO is evaluated using over 700 curated cell culture processes, validated against eight competency questions, and quality-controlled using automated ontology checks. Evaluation results and formal reasoning validation are provided in the Supplementary Materials.

Availability: https://github.com/lewiscelllabs/mcbo

## 1. Introduction

Mammalian cells, such as Chinese Hamster Ovary (CHO) or Human Embryonic Kidney (HEK) cells, are vital in biopharmaceutical manufacturing because they can produce therapeutic proteins, such as monoclonal antibodies, with human-like post-translational modifications, which reduces their immunogenicity in patients[1]. CHO cells are of particular importance. Their ability to be cultured at large industrial scales with high production yields, along with their strong, established history of regulatory approval for their derived products, makes them the “gold standard” for producing biologics[2]. Not only are CHO cells central to biologics manufacturing, they come with decades of accumulated public data[3] and extensive proprietary datasets maintained by industrial partners. These include diverse cell lines, each with proprietary knock-ins or expression cassettes, and rich phenotypic profiles generated under a wide range of cultivation conditions. Yet, despite this data abundance, it remains frustratingly difficult to answer even basic queries (e.g., “*what culture conditions best support a given protein of interest?*”, or “*which mutations improve yield for a particular CHO lineage?*”).

Ontologies provide a structured, hierarchical, and machine-readable representation of a domain, making them a natural solution to these challenges. By formally capturing relationships among cultivation conditions, genetic modifications, and phenotypic outcomes, they can transform scattered datasets into interoperable knowledge that is both human-interpretable and machine-actionable. Beyond this core role, ontologies also complement Large Language Models (LLMs) by adding explicit background knowledge and worldview models to reconcile conflicting perspectives in LLM training data, while enabling tasks such as semantic search and LLM prompt expansion through synonym mappings and domain-specific terminology. Because ontology reasoning is computationally lightweight[4] compared to LLM inference[5] and avoids the overhead of traversals in relational or document-based databases, it provides a practical, cost-efficient foundation for scalable bioprocess data integration[6, 7].

While the broader biomedical community has developed mature ontologies (Gene Ontology[8], ChEBI[9], Cell Line Ontology[10]), and the systems biology community has created sophisticated metabolic models (CHO-GEMs like *i*CHO3K[11]), no comprehensive formal ontology exists yet, specifically for mammalian cell cultivation experimental workflows, for applications in bioprocess development. Existing efforts focus on either broad biological concepts or mathematical modeling frameworks, leaving a gap in practical experimental metadata representation. Recent initiatives show promise, such as NIST’s Industrial Ontology Foundry (IOF [12, 13]) for biopharmaceutical manufacturing, which uses Basic Formal Ontology (BFO[14]) as a foundation for manufacturing process ontologies. However, these efforts have not yet addressed the specific needs of mammalian cell cultivation experimental data capture and integration.

To address this gap, we present a lightweight but extensible mammalian cell cultivation application ontology[15] (AO). Our AO (spoke) is anchored to the IOF (hub) and is authored in *Web Ontology Language* (OWL 2), *Descriptive Logic* (DL)-compatible, so it already supports standard DL reasoners (e.g., HermiT, ELK). While we have not yet added a *Simple Knowledge Organization System* (SKOS) layer, the model is fully compatible with SKOS for synonym and vocabulary management, providing a clear path for standards-compliant expansion. Our goal is not to over-engineer from the start, but to offer a practical first step: a modular, open-source framework that can be deployed internally (e.g., behind firewalls), supports data validation, *Electronic Lab Notebook* (ELN) rendering, and enables progressive enhancement toward *Findable, Accessible, Interoperable, Reusable* (FAIR)[16, 17] data principles and integration with emerging repositories.

## 2. Scope and Competency Questions

We use these competency questions to set the *minimum necessary scope* for MCBO and to guide which upper-ontology commitments and design patterns must be supported. Subsequent sections describe the resulting design choices and how they were implemented and evaluated.

The primary motivation for this ontology is to lay the groundwork for the datahub we are building to support mining of public data. To that end, the scope is limited to the essentials needed to cover the metadata currently captured, and in that way, we avoid over-engineering. More terms will be added as needed, with high ontological commitment, leaving less ambiguity, as is appropriate for an AO. The first release of the datahub will focus on transcriptomic data and key bioprocess parameters, with an emphasis on understanding their interconnections. As we are just now building the datahub, we define rudimentary *Competency Questions* (CQs) for testing MCBO coverage as follows:

**CQ1**: Under what culture conditions (pH, dissolved oxygen, temperature) do the cells (e.g., CHO-K1) reach peak recombinant protein productivity?

**CQ2**: Which CHO cell lines have been engineered to overexpress gene Y?

**CQ3**: Which nutrient concentrations are most associated with viable cell density above Z at day 6 of culture?

**CQ4**: How does the expression of gene X vary between clone A and clone B?

**CQ5**: What pathways are differentially expressed under Fed-batch vs Perfusion in CHO-K1?

**CQ6**: Which are the top genes correlated with recombinant protein productivity in the stationary phase of all experiments?

**CQ7**: Which genes have the highest fold change between cells with viability (>90%) and those without (<50%)?

**CQ8**: Which cell lines or subclones are best suited for glycosylation profiles required for therapeutic protein X?

We further simplify these CQs in the evaluation section to align with the limited metadata currently curated. We will continue to version and share the ontology on GitHub as more data, terms, and CQs are added to support the burgeoning datahub.

## 3. Ontology Design

The *Mammalian Cell Bioprocessing Ontology* (MCBO) is a lightweight application ontology engineered for practical utility in mammalian cell cultivation experiments. It prioritizes semantic correctness, reuse, and extensibility, aligning with Basic Formal Ontology (BFO) and Industrial Ontology Foundry (IOF) standards to support reproducible RNA-seq and culture optimization.

At the heart of the ontology is *CellCultureProcess*, which organizes experimental activities into three primary operational modalities: *BatchCultureProcess* (closed systems), *ContinuousCultureProcess* (continuous medium flow), and *UnknownCultureProcess* (for instances where source metadata is incomplete). Specializations further refine these operational distinctions. *FedBatchCultureProcess* is modeled as a subclass of *BatchCultureProcess* to capture discrete nutrient additions within a closed period. *ContinuousCultureProcess* specializes into *PerfusionCultureProcess* (cell retention) and *ChemostatCultureProcess* (continuous removal of culture broth). Each subclass inherits core relationships, such as participants and outputs, while adding specific operational constraints. Organism specificity is handled through OWL defined classes rather than primitive subclassing. *MammalianCellCultureProcess* is defined as the intersection of *CellCultureProcess* and a participant restriction requiring at least one *MammalianCell*. This supports automated classification and prevents generic bioprocessing templates from being conflated with mammalian-specific logic.

To maintain BFO consistency, MCBO strictly separates occurrents (processes) from continuants (material entities). Culture conditions are not modeled as qualities of the process itself; instead, they inhere in a *CellCultureSystem* (e.g., the bioreactor, medium, and cells) that participates in the process. A *CellCultureProcess RO:has_participant* SOME *CellCultureSystem*, which in turn *RO:has_quality* SOME *CultureEnvironmentalCondition* (e.g., temperature, pH, dissolved oxygen). This preserves the fundamental distinction that processes do not bear qualities. When a condition is actively controlled, an *EnvironmentalSetpoint* (an OBI setting datum) is linked to the quality; the absence of a setpoint indicates the condition was measured but not controlled. Figure 1 illustrates this pattern. Experimental observations are modeled as Information Content Entities (ICE) that are “about” the underlying biological and physical entities. Measurement classes such as *CellViabilityMeasurement, ProductivityMeasurement*, and *GeneExpressionMeasurement* specialize *IAO:0000109* (measurement datum) and carry standardized values and units (e.g., *hasConcentrationValue, hasTiterValue, hasViabilityPercentage*). Similarly, all data products (including *RNASeqDataset, RawReadsDataset*, and *AlignedReadsDataset*) are subclasses of *IAO:dataset*. This ensures that datasets are properly classified as information artifacts rather than material entities or processes. MCBO distinguishes genomic information from manufactured material. A *ProteinProduct* (e.g., a recombinant mAb) is modeled as a material entity linked to its encoding *Gene* via the *encodedByGene* property. This maintains the distinction between the physical output of a bioprocess and the genetic information that specifies it. Gene entities themselves support multiple identifiers, including *hasGeneSymbol* and *hasEnsemblGeneID*, enabling cross-database integration.

**Figure 1.**
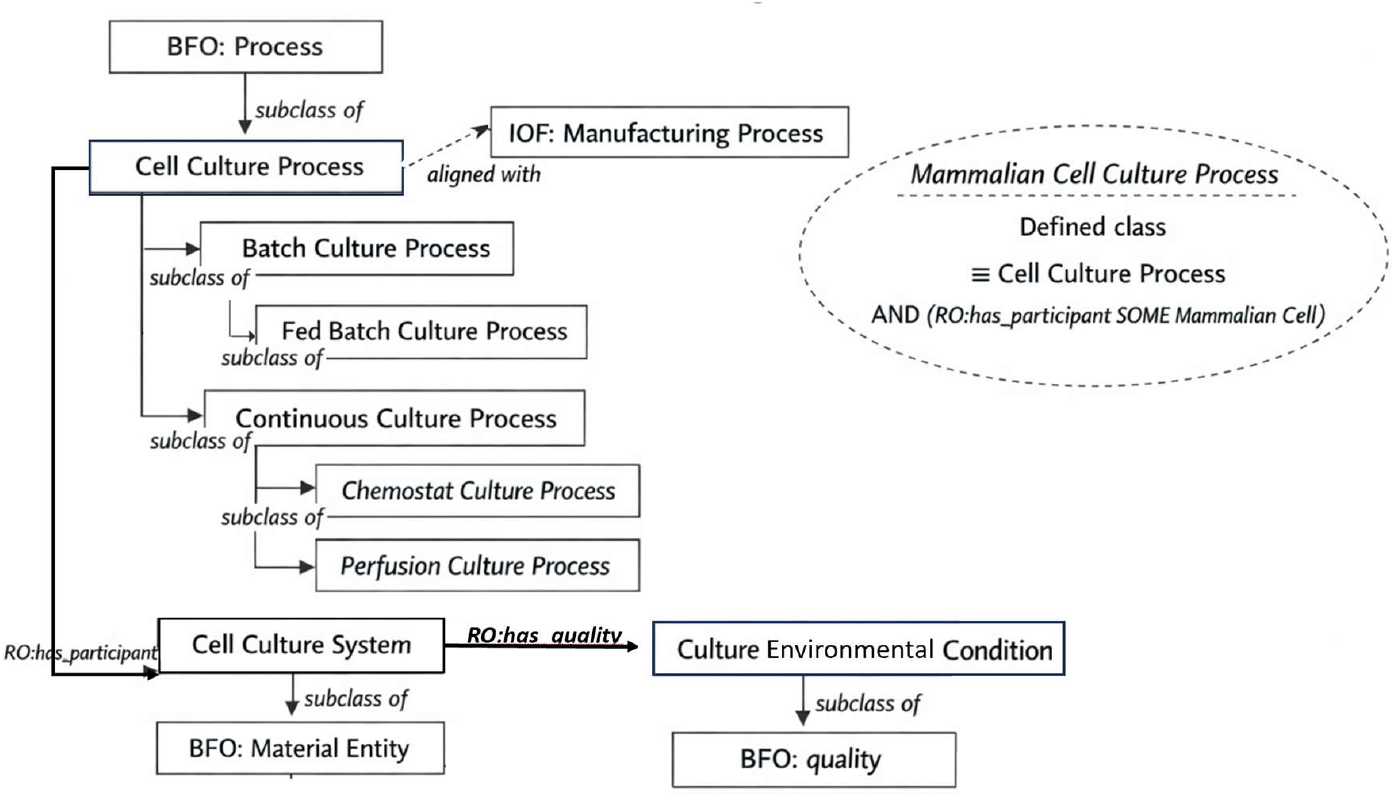
Core process and material entity modeling pattern in MCBO. Generic cell culture processes are organized independently of organism type, with batch and continuous as primary subclasses; fed-batch is modeled as a specialization of batch culture. Mammalian cell culture processes are represented as OWL defined classes using participant restrictions rather than subclassing. Culture environmental conditions are modeled as qualities of the material cell culture system, consistent with BFO principles. Alignment with the IOF Manufacturing Process is shown conceptually without asserting subclass relations.

MCBO adheres to modular reuse, aligning with the Systems Biology Ontology (SBO), Ontology for Biomedical Investigations (OBI), and Units of Measurement Ontology (UO). While the ontology anchors process classes like *CellCultureProcess* to *iof:ManufacturingProcess* via *rdfs:seeAlso* annotations, it maintains direct BFO classification to ensure compatibility with both industrial and biomedical semantic stacks. These alignments provide a pragmatic anchor for downstream data integration and automated reasoning.

## 4. Ontology Development

Our development followed an agile, stakeholder-driven methodology designed to address immediate data integration challenges in the biopharmaceutical manufacturing community. Rather than pursuing comprehensive domain coverage from the outset, we adopted an iterative approach, prioritizing practical utility and extensibility. Initial development was motivated by extensive consultation with industry and academic partners. While the academic community articulated sophisticated use cases, industry partners sought a pragmatic “knowledge base” to bridge the gap between raw data and actionable insights. Through this consultation, a core guiding use case emerged: *“What datasets exist that up-regulate my proteins of interest, and how are they characterized regarding genetic mutations, phenotypes, metabolic states, inputs, and cultivation conditions?”* This question drove our ontological scope as we parse and curate metadata across the vast landscape of publicly available data.

The design was further constrained by several technical imperatives: supporting agile schema evolution as new use cases emerge, enabling auto-generation of data collection interfaces from semantic specifications, and facilitating LLM-assisted extraction of structured metadata from unstructured publications and electronic lab notebooks (ELNs). To meet these goals, ontology was developed using Python-based tooling with a *Terse RDF Triple Language* (TTL) representation, prioritizing integration with modern web technologies and schema validation frameworks. This development cycle leveraged LLM-assisted analysis of existing publications and semi-structured data to identify common terminology and metadata patterns, ensuring the ontology reflects real-world experimental documentation.

## 5. Evaluation

Beyond conceptual design, we evaluated MCBO using real and synthetic datasets to assess coverage, correctness, and practical queryability. Evaluation included ontology quality control checks, population of a real-world knowledge graph derived from published studies, and execution of competency questions implemented as SPARQL queries. Detailed statistics and results are provided in the Supplementary Material, reproducible by publicly available tools, and also summarized here. Detailed evaluation artifacts, including dataset statistics, competency question execution results, and ontology reasoning checks, are provided in the Supplementary Materials.

We assessed the ontology’s ability to represent real-world experimental metadata by selecting three manually annotated cultivation studies from the literature[21–23]. For each study, we identified key metadata (including cultivation strategy, cell line, feeding protocols, measurement types) and verified they could be formally represented using our MCBO terms. Domain experts (JM, NL) reviewed the structure and hierarchical relationships for accuracy relative to current cultivation practices, focusing on whether the ontology captures the essential concepts needed for experimental annotation and data integration.

As of this writing, we have successfully integrated 723 cell culture process instances from published studies[24]. These include 325 unique, real-world bioprocess samples across the integrated runs. The process breakdown includes Batch (518), Fed-batch (135), Perfusion (48), and Unknown (22) operational modalities. All evaluation artifacts, including implementation SPARQL queries and query outputs, are available in the project repository. While curation is ongoing and some real-world datasets currently lack the specific fields (such as detailed gene expression or nutrient concentrations) needed for all 8 competency questions, we supplement the evaluation with a publicly shareable synthetic dataset to demonstrate full functionality.

Real-world data results for the currently populated fields include:

**CQ1 (Culture optimization)**: 161 results correlating culture qualities (pH, dissolved oxygen, temperature) with productivity measurements (*medium* to *high*).

**CQ2 (Cell engineering)**: 3 engineered CHO cell line entries with gene overexpression annotations and their performance metrics.

**CQ5 (Process comparison)**: Successful identification and distribution of all 4 process types across the integrated samples.

All competency questions executed efficiently on the full dataset using consumer-grade hardware, with median query execution times under six (6) seconds for heavy, real-world knowledge graph queries and sub-second performance for the synthetic demonstration data (Supplementary Data S7).. The semantic representation allows for multi-hop graph traversals that harmonize heterogeneous experimental data into a queryable knowledge graph, enabling systematic analysis regardless of the source study structure. This approach demonstrates effective data integration and provides a foundation for cross-study comparative analysis. Full competency question definitions, queries, and outputs are provided in Supplementary Data S3–S5.

To support reviewer evaluation without requiring access to private metadata, we provide a demonstration dataset in data.sample containing synthetic studies designed to exercise all 8 competency questions. Tables 1 and 2 contrast the results between the real curated data and the demonstration data.

**Table 1:**
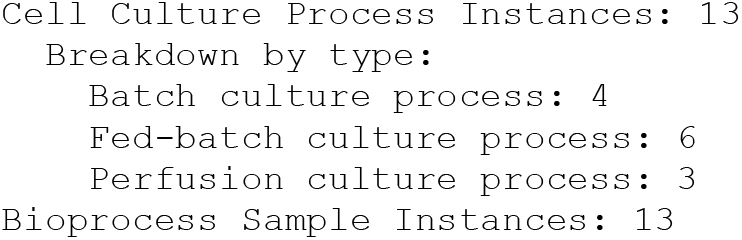
Demo data. statistics (from `mcbo-stats --data-dir data.sample`):

**Table 2:** CQ Results comparison. (from mcbo-run-eval --data-dir [.data|data.sample])

| CQ | Real Data (723 processes) | Demo Data (10 processes) | Notes |
| --- | --- | --- | --- |
| **CQ1** | 161 | 15 | Culture conditions for productivity |
| **CQ2** | 3 | 2 | Overexpression engineering |
| **CQ3** | 0 | 4 | Nutrient concentrations (requires CollectionDay, ViableCellDensity) |
| **CQ4** | 0 | 144 | Gene expression between clones (requires expression matrix) |
| **CQ5** | 4 | 3 | Process type distribution |
| **CQ6** | 0 | 44 | Genes correlated with productivity (requires expression data) |
| **CQ7** | 0 | 7 | Viability fold change (requires ViabilityPercentage) |
| **CQ8** | 0 | 5 | Quality profiles (requires TiterValue, QualityType) |

The demonstration data includes all required fields to show complete functionality. Users can reproduce these results by building the demo graph using the mcbo-build-graph and mcbo-run-eval commands as detailed in the project documentation. Dataset statistics are summarized in Supplementary Data S1. Automated quality control results are provided in Supplementary Data S2.These results demonstrate that MCBO supports cross-study querying of culture conditions, cell engineering strategies, and process modalities using both real and synthetic data, while remaining logically consistent under OWL 2 DL reasoning (Supplementary Data S6). The MCBO ontology evaluated in this study is provided (Supplementary Data S8), enabling independent inspection and reproduction of the reported reasoning results.

## 6. LLM-Powered Query Agent

To demonstrate practical utility beyond manual SPARQL querying, we developed an LLM-powered agent that enables natural language interaction with MCBO-curated data. The agent translates user questions into appropriate SPARQL queries, executes statistical analyses, and synthesizes results into human-readable answers.

The agent follows a tool-calling paradigm where an LLM orchestrator selects from a library of specialized tools: 1) Statistical analysis including correlation, 2) fold-change, and 3) differential expression calculations; Pathway enrichment via KEGG[25] and Reactome[26] API integration. The agent supports multiple LLM backends (OpenAI GPT-4, Anthropic Claude, and local models via Ollama), enabling deployment in both cloud and privacy-sensitive on-premises environments.

### Example interaction

~~~
**User**: “What genes are differentially expressed under Fed-batch
    vs Perfusion in HEK293?”
**Agent**: [Executes SPARQL template for process-type comparison]
       [Computes fold-change and statistical significance]
       [Returns ranked gene list with p-values]
~~~

The agent implements the Model Context Protocol (MCP), enabling integration with AI assistants and other LLM-aware ecosystems. As an MCP server, the query agent allows researchers to query MCBO data directly from their development environment without writing SPARQL. The agent is also available as a command-line tool (‘mcbo-agent-eval‘). Full documentation is provided at ‘docs/agent.md‘ in the repository.

Agent Documentation: https://mcbo.readthedocs.io/en/latest/agent.html

## 7. Discussion

The MCBO ontology addresses a critical gap in biopharmaceutical manufacturing: the lack of a standardized, structured, metadata framework that is both biologically specific and industrially interoperable. By prioritizing the development of a formal ontology before defining a rigid data schema, we have created a foundation that is independent of specific database implementations.. This separation of structure from content allows the ontology to serve as a stable substrate for harmonizing heterogeneous data across public repositories and proprietary internal systems. Formal logical consistency and reasoning-based validation of the ontology are documented in Supplementary Section S6.

A key advantage of this layered approach is its compatibility with industrial security requirements. Because MCBO is open-source and uses standard web technologies, it can be deployed behind institutional firewalls. This enables partners to integrate public benchmarks with private experimental data for federated querying and foundation model refinement without exposing sensitive intellectual property. By filling the gaps in existing vocabularies like SBO and OBI with practical, cultivation-specific terms and constraints, MCBO transforms ad hoc data curation into a structured, actionable knowledge base.

This semantic foundation enables a shift toward agentic and human-in-the-loop workflows. Rather than requiring users to write complex code, the infrastructure supports natural language interaction where AI agents execute semantic queries, subset datasets, and generate visualizations (e.g., “Analyze gene expression in Fed-batch cultures of CHO-K1 with glucose limitation and high IgG yield”). To bootstrap this system, we currently leverage LLM-powered pipelines that extract structured metadata from legacy documents (e.g., ELNs, publications). This technology stack automates generation of provenance-aware records and pre-populated dataset submission forms, reducing friction and increasing consistency across submissions to our datahub. The MCBO-schema framework serves as a scalable substrate for building intelligent, user-friendly, and future-proof mammalian bioprocessing data infrastructures. It paves the way for a transition from fragmented data silos to an AI-native ecosystem, grounding innovation in principled semantic foundations.

## 8. Future work

Future iterations of MCBO will focus on deepening industrial integration and expanding the scope of supported datasets. Future development will prioritize alignment with established initiatives, particularly the Industrial Ontology Foundry (IOF) biopharmaceutical manufacturing extensions and their underlying Basic Formal Ontology (BFO)[28] foundations. We will continue to systematically map our cultivation concepts to IOF Core constructs and we aim to align our modeling pathways with industry standards from bodies such as the *BioPhorum Operations Group* (BPOG)[29], *Bio-Process Systems Alliance* (BPSA), and *International Society for Pharmaceutical Engineering* (ISPE). To strengthen semantic rigor, we will formalize cross-mappings to established biomedical standards, including: SBO, OBI, CLO, *Minimum Information About a Proteomics Experiment-MIAPE*[30], *Minimum Information about a High-Throughput Nucleotide Sequencing Experiment* - MINSEQE[31], *Minimum Information Required for A Glycomics Experiment* - MIRAGE[32], *Metabolomics Standards Initiative* - MSI[33], *Investigation–Study–Assay Tab-delimited format* - ISA-Tab[34], *Minimum Information for Biological and Biomedical Investigations* - MIBBI[35], *Minimum Information Required in the Annotation of Models* - MIRIAM[36], *Minimum Information About a Simulation Experiment* - MIASE[37].

A primary near-term goal is the exploration of semantic bridges to existing CHO Genome-Scale Metabolic Models (CHO-GEM)[27]. By providing a formal ontological grounding for metabolic model annotations, we can enable integrated queries that correlate real-world cultivation conditions with predicted metabolic flux. This will be supported by comprehensive competency question validation as we integrate larger mammalian bioprocessing multi-omics datasets into the datahub. SKOS definitions will be developed to strengthen semantic rigor. Near-term priorities include comprehensive competency question validation using real mammalian bioprocessing omics datasets, formal evaluation against existing literature, and expert review by industry practitioners, all driven by real use cases as we grow our datahub.

MCBO is committed to an open, community-driven development model. All ontology files, schema definitions, and supporting tools are available under the permissive MIT license. We establish formal feedback loops with academic labs, biofoundries, and industrial partners through specialized GitHub issue templates and review processes. Longer-term, we intend to register MCBO with BioPortal and identifiers.org, pursue OBO Foundry acceptance, and coordinate closely with relevant standards bodies to ensure the ontology evolves alongside the needs of the biopharmaceutical manufacturing ecosystem.

## Supporting information

Dataset Scale and Composition

Ontology Quality Control

Competency Questions and Query Results

Real-World Data Evaluation Results

Synthetic Demonstration Dataset Results

Ontology Reasoning and Logical Consistency

Query Performance Benchmark

The MCBO Ontology File

## Acknowledgements

NEL, KR were supported in part by funding from the National Institutes of Health (R35 GM119850), and support from the Georgia Research Alliance. iBioNe is supported by National Science Foundation Award #2114716.

## Glossary of Acronyms

AI: Artificial Intelligence
AO: Application Ontology
BFO: Basic Formal Ontology
BPOG: BioPhorum Operations Group
BPSA: Bio-Process Systems Alliance
CHO: Chinese Hamster Ovary (cells)
ChEBI: Chemical Entities of Biological Interest
CLO: Cell Line Ontology
CQ: Competency Question
ELN: Electronic Laboratory Notebook
FAIR: Findable, Accessible, Interoperable, Reusable
GEM: Genome-scale Metabolic Model
GPT: Generative Pre-trained Transformer
IAO: Information Artifact Ontology
ICE: Inventory of Composable Elements
IOF: Industry Ontology Foundry
IP: Intellectual Property
ISPE: International Society for Pharmaceutical Engineering
KEGG: Kyoto Encyclopedia of Genes and Genomes
LLM: Large Language Model
MCBO: Mammalian Cell Bioprocessing Ontology
MCP: Model Context Protocol
MIAPE: Minimum Information About a Proteomics Experiment
MIT: Massachusetts Institute of Technology
OBI: Ontology for Biomedical Investigations
OBO: Open Biological and Biomedical Ontologies
OWL: Web Ontology Language
pH: Potential of Hydrogen
RDF: Resource Description Framework
RNA: Ribonucleic Acid
SBO: Systems Biology Ontology
SKOS: Simple Knowledge Organization System
SPARQL: SPARQL Protocol and RDF Query Language
UO: Units Ontology

## Supplmentary Materials

The supplementary materials provide quantitative evaluation of MCBO, including dataset scale statistics (S1), ontology quality control (S2), competency question implementation and execution (S3–S5), and formal reasoning-based consistency verification (S6), and SPARQL query performance benchmarks (S7), and the MCBO ontology file itself (Supplementary Data S8).

