## Supplementary material for "MCBO: Mammalian Cell Bioprocessing Ontology, A Hub-and-Spoke, IOF-Anchored Application Ontology": Ontology Reasoning and Logical Consistency

### S6 Protégé Reasoning Results

Logical consistency of the MCBO ontology was evaluated using the HermiT OWL 2 DL reasoner (version 1.4.3.456) within Protégé. The ontology and all declared imports loaded successfully under reasoning, indicating that the ontology is syntactically well-formed and that all referenced external ontologies are resolvable. Figure S6-1 summarizes ontology metadata, axiom statistics, and resolved imports.

The ontology was then classified using HermiT. Classification completed without errors and produced an inferred class hierarchy containing no unsatisfiable named classes, indicating the absence of internal logical contradictions (Figure S6-2).

As an additional model-existence sanity check, a DL query was executed for the contradictory class expression *owl:Thing* $\sqcap \neg$*owl:Thing*. The reasoner correctly inferred this expression to be unsatisfiable, confirming that the ontology admits at least one model and is therefore logically consistent under OWL 2 DL semantics (Figure S6-3).

**Figure S6-1**: MCBO ontology loaded in Protégé, showing ontology metadata, axiom statistics, and resolved imports.

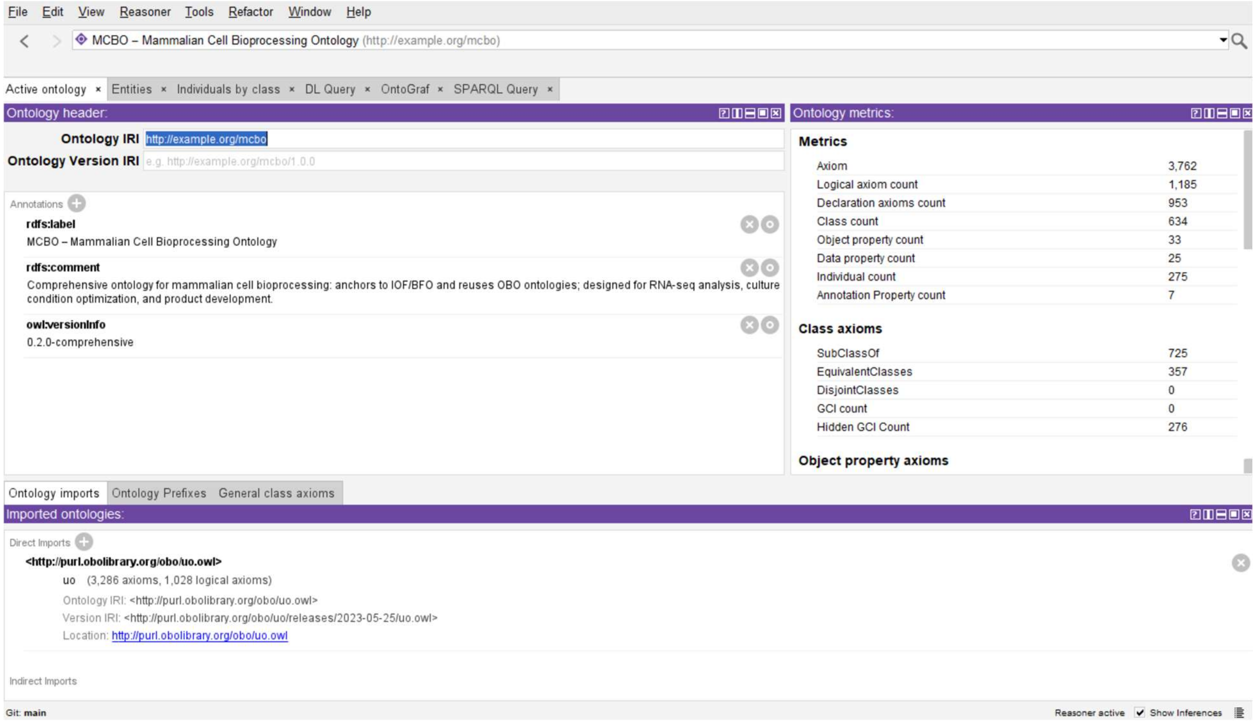

**Figure S6-2.** Inferred class hierarchy of the MCBO ontology after classification with the HermiT OWL 2 DL reasoner in Protégé, showing no unsatisfiable named classes.

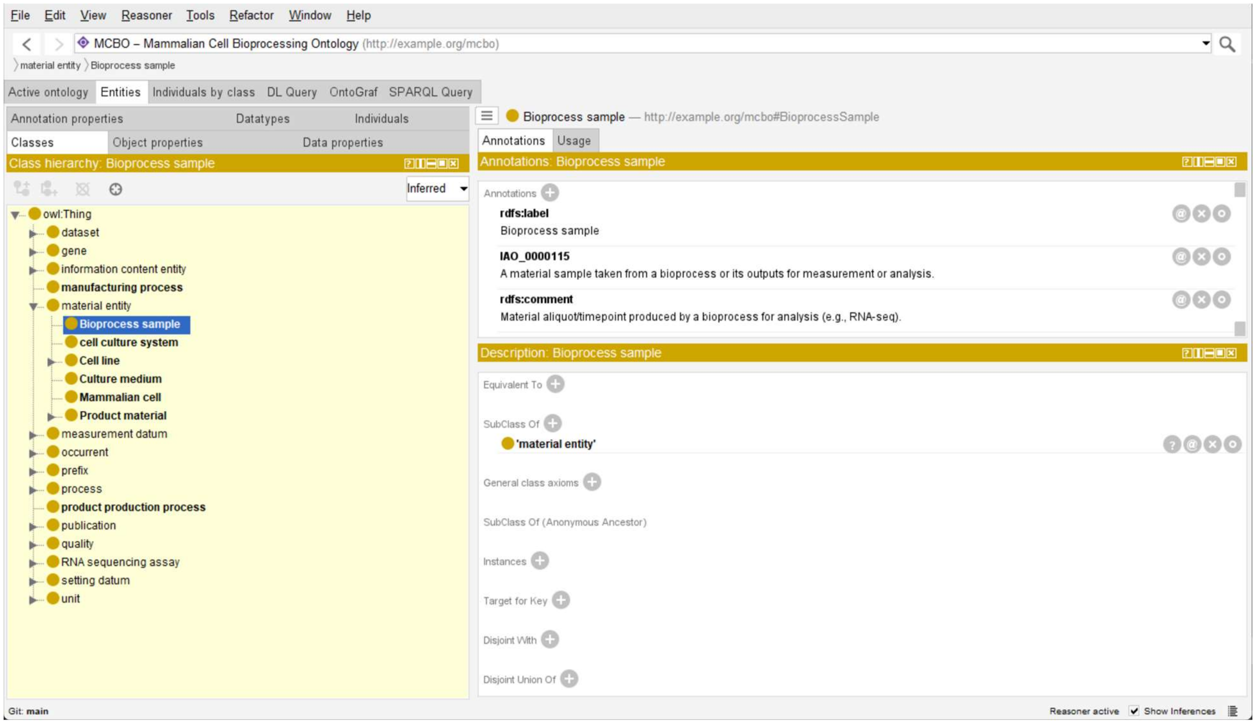

**Figure S6-3**: DL query in Protégé demonstrating the unsatisfiability of the contradictory class expression *owl:Thing* ⊓ ¬*owl:Thing*, confirming logical consistency of the MCBO ontology under HermiT reasoning.

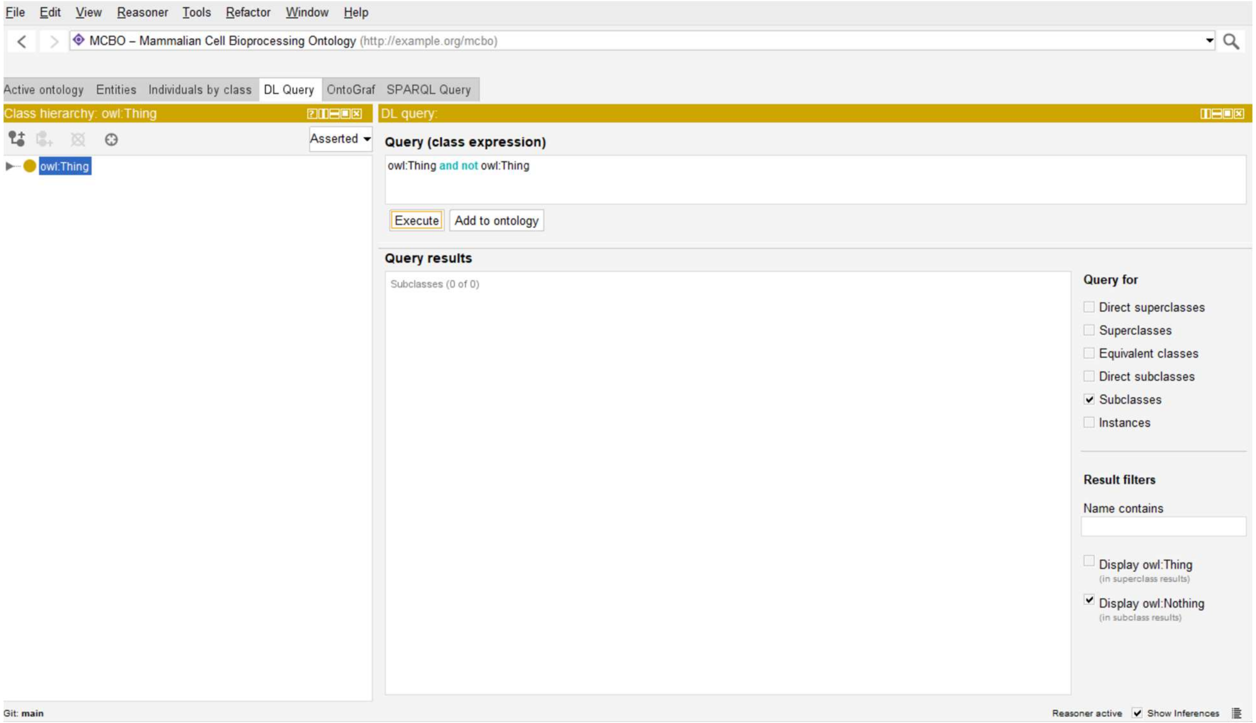
