## Supplementary material for "MCBO: Mammalian Cell Bioprocessing Ontology, A Hub-and-Spoke, IOF-Anchored Application Ontology": Query Performance Benchmark

### S7 Query Performance Benchmark

To assess the operational performance of MCBO in realistic usage scenarios, we measured execution times for all eight competency questions implemented as SPARQL queries over both the real-world and synthetic knowledge graphs. The real-world graph comprises 723 instantiated cell culture processes derived from curated source data, while the synthetic graph represents a minimal demonstration dataset containing 10 processes with complete coverage of all competency questions.

Query execution times were measured under a single-user, local execution setting to reflect typical interactive analysis workflows rather than optimized high-performance deployments. Table S7-1 reports observed execution times for each competency question on both datasets. While query complexity and result cardinality varied across questions, all queries completed successfully, with most executing in sub-second time on the real-world dataset where data were limited, and all completing in under seven seconds. Queries executed against the much smaller, synthetic dataset consistently completed in under one second.

These results demonstrate that MCBO supports practical, executable querying over hundreds of instantiated bioprocesses using standard SPARQL tooling on commodity hardware, providing sufficient performance for exploratory analysis and iterative knowledge graph interrogation.

**Execution environment.** All benchmarks were conducted on a local workstation running Linux (WSL2) with an 11th-generation Intel Core i7-11800H CPU (16 cores) and 31.2 GB RAM, using Python 3.10.19. Detailse

**Table S7-1**: SPARQL Query Execution Time

| Competency Question | Real Data (723 processes) | Synthetic Data (10 processes) |
|---|---|---|
| CQ1 | 6.80 s | 0.20 s |
| CQ2 | 5.50 s | 0.00 s |
| CQ3 | 0.10 s | 0.10 s |
| CQ4 | 0.10 s | 0.50 s |
| CQ5 | 0.10 s | 0.00 s |
| CQ6 | 0.30 s | 0.20 s |
| CQ7 | 0.10 s | 0.50 s |
| CQ8 | 0.30 s | 0.10 s |
